# Neuroinvasion and neurotropism by SARS-CoV-2 variants in the K18-hACE2 mouse

**DOI:** 10.1101/2021.04.16.440173

**Authors:** Frauke Seehusen, Jordan J. Clark, Parul Sharma, Eleanor G Bentley, Adam Kirby, Krishanthi Subramaniam, Sabina Wunderlin Giuliani, Grant L. Hughes, Edward I. Patterson, Benedict D. Michael, Andrew Owen, Julian A. Hiscox, James P. Stewart, Anja Kipar

## Abstract

Severe Acute Respiratory Syndrome Coronavirus 2 (SARS-CoV-2) not only affects the respiratory tract but also causes neurological symptoms such as loss of smell and taste, headache, fatigue or severe cerebrovascular complications. Using transgenic mice expressing human angiotensin-converting enzyme 2 (hACE2) we investigated the spatiotemporal distribution and pathomorphological features in the CNS following intranasal infection with SARS-CoV-2 variants, also after prior influenza A virus infection. Apart from Omicron, we found all variants to frequently spread to and within the CNS. Infection was restricted to neurons and appeared to spread from the olfactory bulb mainly in basally orientated regions in the brain and into the spinal cord, independent of ACE2 expression and without evidence of neuronal cell death, axonal damage or demyelination. However, microglial activation, microgliosis and a mild macrophage and T cell dominated inflammatory response was consistently observed, accompanied by apoptotic death of endothelial, microglial and immune cells, without their apparent infection. Microgliosis and immune cell apoptosis indicate a potential role of microglia for pathogenesis and viral effect in COVID-19 and possible impairment of neurological functions, especially in long COVID. These data may also be informative for the selection of therapeutic candidates, and broadly support investigation of agents with adequate penetration into relevant regions of the CNS.

## Introduction

As of March 2022 the newly emerged betacoronavirus, Severe Acute Respiratory Syndrome Coronavirus-2 (SARS-CoV-2) has infected over 476 million people globally, with over 6.1 million deaths due to the associated disease, COVID-19 (WHO, COVID-19 Dashboard, 27th March 2022). The majority of COVID-19 patients display respiratory symptoms. However, a proportion of patients develop mostly transient unspecific neurological signs like loss of smell and taste (anosmia, ageusia), headache, or dizziness. Fatal cases can also be associated with ischemic stroke, hemorrhagic encephalopathy and epileptic seizures as well as meningoencephalitis [1–7].

There is evidence of viral entry into the brain through the olfactory or vagal nerve and/or the oral and ophthalmic routes, with transsynaptic neuronal spread to other brain regions [8–10]; this could provide a link to the frequently reported (initial) anosmia and ageusia. However, sparing of the olfactory bulb despite infection of olfactory and nasal mucosa was also detected [11]. In addition, hematogenous spread to the brain via infection of endothelial cells and/or immune cells has been suspected, though since been disputed [12–14].

Pathomorphological findings reported in the brain of fatal human COVID-19 cases are variable and present as vascular/hemodynamic/ischemic lesions like ischemic infarcts and/or mild inflammatory changes; neuronal or axonal damage and acute disseminated encephalomyelitis have been reported in rare cases [9,15–22]. Focal or diffuse microglial activation or microglial nodules have also been observed [9,15- 17,20,23–24]. COVID-19 patients can harbor SARS-CoV-2 RNA and protein in the brain [9-10,25], and a study using human brain organoids provided strong evidence that SARS-CoV-2 replicates in neurons [26]. However, presence of the virus was not found to be associated with the severity of neuropathological changes [9], and a report on 18 patients failed to detect SARS-CoV-2 antigen by immunohistochemical staining although low levels of viral RNA were detected by RT-PCR in five patients [21]. Another study found viral RNA in the leptomeningeal layer surrounding the olfactory bulb and interpreted this finding as evidence against the potential neurotropic properties and neuroinvasive capacity of SARS-CoV-2 [11].

Encephalopathy, as a severe complication in COVID-19, is often associated with systemic hyperinflammation mainly provoked by an aberrantly excessive innate immune response [27]. It is suspected that not only a direct virus induced endotheliitis, but also a maladaptive innate immune response may impair neurovascular endothelial function and cause disruption of the blood-brain barrier (BBB), activation of innate immune signal-ling pathways and a parainfectious autoimmunity [27]. However, infection of endothelial cells has been questioned subsequently [14], thereby still awaiting final proof or refute.

The role of glial cells in COVID-19 encephalopathies has also been discussed [28–29]. Glial cells might not only represent potential targets for viral infection but are also highly sensitive to systemic proinflammatory cytokines [30–32]. In COVID-19 patients, the massive systemic release of inflammatory cytokines could affect endothelial cells and astrocytes of the BBB, thus facilitating viral entry like in other viral brain diseases such as HIV-1 encephalitis and measles [33]. Indeed, it has been shown that SARS-CoV-2 spike protein S1 from the blood can pass the BBB and thereby gain access to the brain parenchyma in mice, potentially triggering a parenchymal response without the presence of intact virus [34]. Furthermore, astrocytes and microglia may contribute to the local neuroinflammatory response of the CNS [28, 35]. SARS-CoV-2 infection of astrocytes has been described in human brains [36]. Also, microglial cells are hypothesized to be involved in the innate immune response and facilitate viral clearance, recruitment of immune cells as well as activation of antiviral responses and cytokine production in the brain of COVID-19 patients [29]. Some authors also propose that a SARS-CoV-2-induced proinflammatory microglial phenotype might contribute to the development of subsequent neurodegenerative disorders [37–38]. Furthermore, the proinflammatory priming of microglia, either by direct SARS-CoV-2 infection or a peripheral cytokine storm, could exacerbate disease, as indicated in experimental murine coronavirus (MHV-A59) infection [39].

The human angiotensin-converting enzyme 2 (hACE2) is considered to be the main host receptor for SARS-CoV-2, binding to the viral spike protein (S) [40]. The K18-hACE2 transgenic (K18-hACE2) mouse, where hACE2 expression is driven by the epithelial cell cytokeratin-18 (K18) promoter, developed to study the pathogenesis of SARS-CoV infection [41], is frequently employed to address a broad range of questions regarding SARS-CoV-2 as expression of hACE2 appears to convey higher binding affinity than its murine counterpart. Several studies have shown that intranasal SARS-CoV-2 infection of K18-hACE2 mice reaches the brain where it spreads widely. Infection of the brain can be associated with non-suppurative (meningo)encephalitis [26,42–44]. With high infectious doses (105 PFU), very high viral loads were found in the brain at a time when lung burdens had already decreased, in association with upregulation of IFN-α as well as proinflammatory cytokine and chemokine transcription; the authors proposed that clinical symptoms or lethal outcome of infection in these mice was a consequence of neuroinvasion [44–45]. Another study using an even higher viral inoculum (106 PFU) elicited CNS signs (tremors, proprioceptive defects, abnormal gait, imbalance) by day 6/7 after infection [42]. Studies in mice have also confirmed that the virus can get access to the brain via the olfactory bulb [26, 34].

Detailed information on the effect of SARS-CoV-2 in the brain of COVID-19 patients has so far been collected from fatal cases. However, there is a current paucity of data relating to what happens in the brain of patients that do not have severe disease, recover from it or develop long COVID. A better understanding of the viral dynamics within the CNS will provide further insight into the disease pathogenesis and will be highly informative for therapeutics development by providing insight into the prerequisites for distribution of new medicines in development. The present study represents an attempt to address this question by investigating the CNS of K18-hACE2 mice after intranasal infection with low to moderate doses of SARS-CoV-2 strains, also in combination with prior influenza A virus infection as it might foster neuronal spread [43].

## Material and Methods

### Cell culture and virus

A Pango lineage B strain of SARS-CoV-2 (hCoV-2/human/Liverpool/REMRQ0001/ 2020) cultured from a nasopharyngeal swab from a patient, was passaged in Vero E6 cells [46]. The fourth virus passage (P4) was used for infections after it had been checked for deletions in the mapped reads and the stock confirmed to not contain any deletions that can occur on passage [43].

Human nCoV19 isolate/England/202012/01B (lineage B.1.1.7; Alpha variant) was from the National Infection Service at Public Health England, Porton Down, UK via the European Virus Archive (catalogue code 004V-04032). This was supported by the European Virus Archive GLOBAL (EVA-GLOBAL) project that has received funding from the European Union’s Horizon 2020 research and innovation programme under grant agreement No 871029.

B.1.351 (Beta variant: 20I/501.V2.HV001) isolate [47] was received at P3 from the Cen-tre for the AIDS Programme of Research in South Africa (CAPRISA), Durban, in Oxford in January 2021, passaged in VeroE6/TMPRSS2 cells (NIBSC reference 100978), used here at P4. Identity was confirmed by deep sequencing at the Wellcome Trust Centre for Human Genetics, University of Oxford.

The B.1.617.2 (Delta variant) hCoV-19/England/SHEF-10E8F3B/2021 (GISAID acces-sion number EPI_ISL_1731019) was kindly provided by Prof. Wendy Barclay, Imperial College London, London, UK through the Genotype-to-Phenotype National Virology Consortium (G2P-UK). Sequencing confirmed it contained the spike protein mutations T19R, K77R, G142D, Δ156-157/R158G, A222V, L452R, T478K, D614G, P681R, D950N. The B.1.1.529/BA.1 (Omicron variant) isolate M21021166 was originally isolated by Prof Gavin Screaton, University of Oxford, UK [48] and then obtained from Prof. Wendy Barclay, Im-perial College London, London, UK through G2P-UK. Sequencing confirmed it contained the spike protein mutations A67V, Δ69- 70, T95I, G142D/Δ143-145, Δ211/L212I, ins214EPE, G339D, S371L, S373P, S375F, K417N, N440K, G446S, S477N, T478K, E484A, Q493R, G496S, Q498R, N501Y, Y505H, T547K, D614G, H655Y, N679K, P681H, N764K, A701V, D796Y, N856K, Q954H, N969K, L981F.

### The titres of all isolates were confirmed on Vero E6 cells and the sequences of all stocks confirmed

Influenza virus A/HKx31 (X31, H3N2) was propagated in the allantoic cavity of 9- day-old embryonated chicken eggs at 35 °C. Titres were determined by an influenza plaque assay using MDCK cells [43].

### Biosafety

All work was performed in accordance with risk assessments and standard operat-ing procedures approved by the University of Liverpool Biohazards Sub-Committee and by the UK Health and Safety Executive. Work with SARS-CoV-2 was performed at containment level 3 by personnel equipped with respirator airstream units with filtered air supply.

### Animals and virus infections

Animal work was approved by the local University of Liverpool Animal Welfare and Ethical Review Body and performed under UK Home Office Project Licence PP4715265. Mice carrying the human ACE2 gene under the control of the keratin 18 promoter (K18-hACE2; formally B6.Cg-Tg(K18-ACE2)2Prlmn/J) were purchased from Jackson Laboratories and Charles River. Mice were maintained under SPF barrier conditions in individually ventilated cages.

For each experiment, animals were randomly assigned into multiple groups. For SARS-CoV-2 infection, mice were anaesthetized lightly with isoflurane and inoculated in-tra-nasally with 50 µl containing 103 PFU or 104 PFU (cohort 3; Pango lineage B) SARS-CoV-2 in PBS (Table 1); control animals received PBS. For double infections (cohort 2), mice were anaesthetized lightly with KETASET i.m. and inoculated intranasally with 102 PFU IAV X31 in 50 µl sterile PBS. Three days later, they were infected with SARS-CoV-2 (Pango lineage B), as described above. Mock-infected mice served as controls.

**Table 1.**
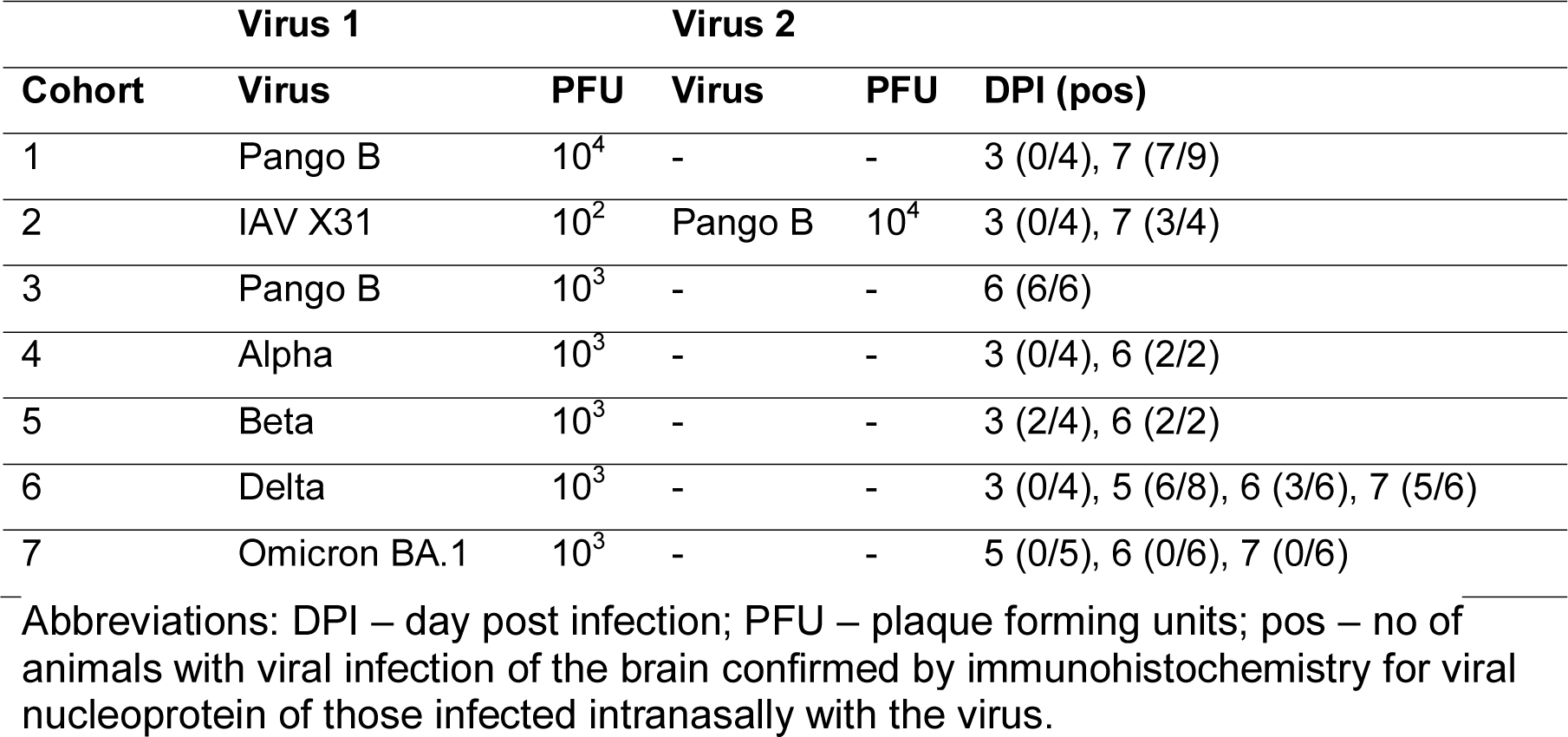
Study cohorts. All viruses, unless otherwise stated, are SARS-CoV-2 strains. Animal num-bers differ, as for some cohorts, mice originated from several experiments. For SARS-CoV-2 Alpha and Beta, only a few animals were examined, to determine whether the strain can also spread to the brain. In Cohort 3, SARS-CoV- 2 Pango lineage B infection took place 3 days after IAV infection.

Mice were monitored for any clinical signs and weighed. Animals were sacrificed at 3, 5, 6 or 7 days post SARS-CoV-2 infection (Table 1) by an overdose of pentabarbitone. Mice were dissected and tissues collected immediately for downstream processing; the lungs were processed for other studies [43,49–50].

### Tissue collection, preparation and processing

The heads from all animals and the spinal cords (C1-T12) of selected animals of cohorts 1 and 2 were collected and fixed in 10% neutral buffered formal saline for 24-48 h. For cohorts 1, 2, 4, 5 and part of cohort 6, brains were exenterated and coronal sections prepared. Heads were then sawn longitudinally in the midline using a diamond saw (Exakt 300; Exakt) for histological assessment of the nasal cavity, cribriform plate and rostral parts of the olfactory bulb that had not been removed with the brain. For all other animals, heads were sawn longitudinally in the midline and the brain left in the skull. In cases from cohorts 1 and 2 where SARS-CoV-2 viral antigen was detected in the brain (see below), the spinal cords were sawn into appr. 1.5 mm thick cross sections. Heads and spinal cord sections were gently decalcified in RDF (Biosystems) for twice 5 days, at room temperature (RT) and on a shaker. Brains, heads and spinal cords were routinely paraffin wax embedded.

### Histology, immunohistology

Consecutive sections (3-5 µm) were either stained with hematoxylin and eosin (HE) or used for immunohistochemistry (IH). IH was performed using the horseradish peroxidase (HRP) method to detect viral antigen in all examined tissues in all animals, and in selected cases to identify macrophages/activated microglial cells (Iba1+), T cells (CD3+), B cells (CD45R/B220+) and neutrophils (Ly6G+) and to highlight astrocytes (glial fibrillary acidic protein, GFAP+), apoptotic cells (cleaved caspase 3+), disturbances of the fast axonal transport indicating acutely damaged neuronal axons (amyloid precursor protein, APP+) and ACE2 expression in selected animals where viral antigen was detected in the brains. Antibodies and detection systems are listed in Supplemental Table S1. Briefly, after deparaffination, sections underwent antigen retrieval in citrate buffer (pH 6.0) or Tris/EDTA buffer (pH 9) for 20 min at 98 °C, followed by incubation with the primary antibodies (diluted in dilution buffer, Agilent Dako). This was followed by blocking of endogenous peroxidase (peroxidase block, Agilent Dako) for 10 min at RT and incubation with the appropriate secondary antibodies/detection systems, all in an autostainer (Dako Agilent or Ventana). Sections were subsequently counterstained with hematoxylin.

The brain of a mock-infected control hACE mouse served as normal brain control for ACE2, GFAP and APP, and a lymph node from a normal mouse for the leukocyte and apoptosis markers. Sections from the lungs served as internal positive controls for SARS-CoV-2 and IAV antigen expression, and sections incubated without the primary antibodies served as negative controls.

### RNA in situ hybridization

In selected cases, RNA ISH was performed using the RNAscope® ISH method (Advanced Cell Diagnostics (ACD Advanced Cell Diagnostics, Newark, California) and the RNAscope® 2.5 Detection Reagent Kit (Brown) according to the manufacturer’s protocol and as previously described [51–52]. All cases were first tested for the suitability of the tissue (RNA preservation and quality) with an oligoprobe for Mus musculus peptidylprolyl isomerase B (PPIB) mRNA (ACD). Those yielding good PPIB signals were then subjected to RNA-ISH for nCoV2019-S (coding for Wuhan seafood market pneumonia virus isolate Wuhan-Hu-1 complete genome; Genbank NC_045512.2). Briefly, sections were heated to 60 °C for 1 h and subsequently deparaffinized. Permeabilization was achieved by incubating the section in pretreatment solution 1 (RNAscope® Hydrogen Peroxide) for 10 min at RT, followed by boiling in RNAscope® 1X Target Retrieval Reagents solution at 100 °C for 15 min and washing in distilled water and ethanol. After digestion with RNAscope® Protease Plus for 30 min at 40 °C, sections were hybridized with the oligoprobes at 40 °C in a humidity control tray for 2 h (HybEZTM Oven, ACD).

Thereafter a serial amplification with different amplifying solutions (AMP1, AMP2, AMP3, AMP4: alternating 15 min and 30 min at 40 °C) was performed. Between each incubation step, slides were washed with washing buffer. They were subsequently incubated with AMP 5, AMP 6 and DAB at RT for 30 and 15 min respectively. Gill’s hematoxylin served to counterstain the sections which were then dehydrated with graded alcohol and xylene and coverslipped. A lung section from an infected mouse at 3 dpi served as a positive control. The negative control was consecutive sections incubated accordingly but without including the hybridization step.

## Results

### In K18-hACE2 mice, SARS-CoV-2 Pango B, alpha, beta and delta variant infection is frequently accompanied by virus spread into the central nervous system

To assess whether SARS-CoV-2 gains access to the CNS after intranasal infection with a low to medium viral dose we examined groups of K18-hACE2 mice that had been infected with SARS-CoV-2 Pango lineage B (a variant from the initial outbreak in the UK, strain hCoV-19/England/Liverpool_REMRQ0001/2020 [43], at 103 and 104 PFU; an Alpha variant (B.1.1.7), a Beta variant (B.1.351), a Delta variant (B1.617.2) and a near clinical (B.1.1.529) Omicron variant isolate from the UK [48] (all at 103 PFU). In addition, we examined mice that had been infected with Pango lineage B inoculum (104 PFU) at day 3 post intranasal infection with 102 PFU IAV (strain A/X31). SARS-CoV-2 infected mice began to lose weight at 3 or 4 dpi and continued to lose weight until the end of the experiment, at 5, 6 or 7 dpi, with the exception of the Omicron infected animals which recovered from the weight loss by day 6 (Supplemental Fig. S1) [43, 49–50]. In the sequential IAV then Pango lineage B infected mice as well as some delta variant infected mice, the weight loss was accelerated leading to more rapid and higher mortality, as determined by a humane endpoint of 20% weight loss [43].

At day 3 post intranasal Pango lineage B, Alpha, Beta and Delta variant infection, viral antigen was detected in numerous individual and aggregates of occasionally degenerate epithelial cells in the nasal cavity in all SARS-CoV-2 infected animals and in 2 of the 4 IAV and Pango lineage B double infected animals. Viral antigen was also found in the lungs, in both type I and II pneumocytes in randomly distributed and variably sized patches of alveoli, with only occasional degenerate cells. This was associated with a mild increase in interstitial cellularity, endothelial cell activation with rolling, emigration and perivascular aggregation of some lymphocytes, and small macrophage aggregates, all as previously reported by this group [43]. In dual infections, identical SARS-CoV-2 associated lesions were present within areas unaffected by IAV lesions. At this stage, SARS-CoV-2 antigen was occasionally detected in the olfactory epithelium. It was generally not observed in brain nerves and brain including olfactory bulb; however, the two Beta variant infected mice exhibited several positive neurons in the olfactory bulb and patches of positive neurons in the frontal cortex and brain stem, respectively (Fig. 1). In dual infected animals, IAV antigen expression was not detected in these structures either, and there were no histological changes.

**Figure 1.**
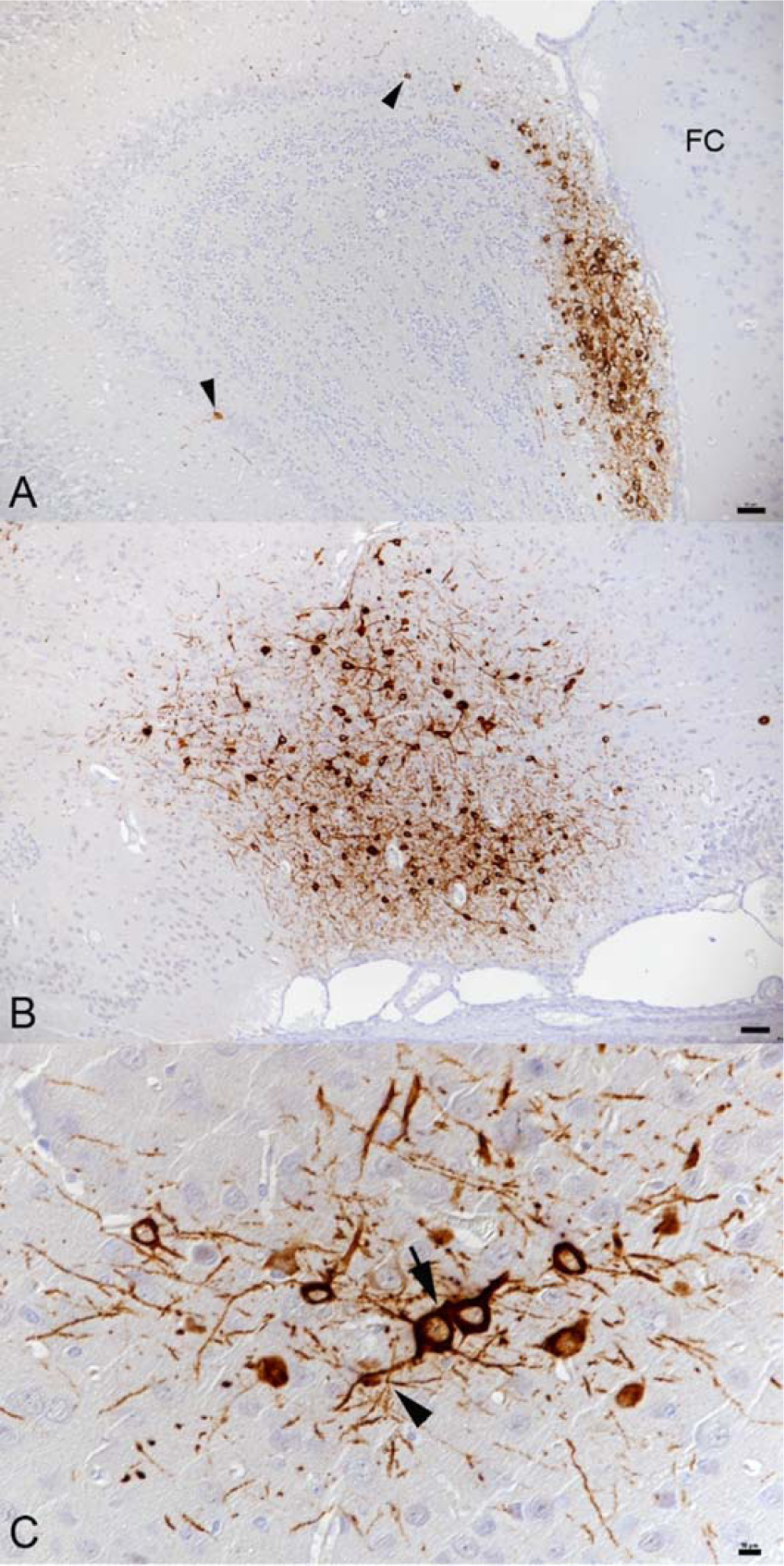
Brain, K18-hACE mice, day 3 post intranasal infection with SARS-CoV-2 Beta variant at 10^3^ PFU. **A)** Olfactory bulb. Viral antigen is detected in a large patch of neurons in granule layer, inner and outer plexiform layer, mitral layer and glomerular layer and scattered individual neurons. FC – frontal cortex. Bar = 50 µm. **B)** Brain stem (anterior olfactory nucleus). Large patch of positive neurons. Bar = 50 µm. **C)** Cortex, same animal as in B. Patch of positive neurons with viral antigen expression in both cell body and processes. Neurons (arrow) and neuronal processes (arrowhead) often appear close apposed. Bar = 10 µm. Immunohistochemistry, hematoxylin counterstain.

At days 5, 6 and 7 post infection, the respiratory and olfactory epithelium of the nasal mucosa still exhibited viral antigen expression, more extensive in Pango lineage B, Alpha, Beta and Delta variant than in Omicron infected mice, as confirmed by differences in viral loads detected in throat swabs [49], and decreasing with time. Infected cells generally appeared unaltered. At day 5 (studied in Delta and Omicron infected mice) the lungs of Delta variant infected mice exhibited a mild to moderate multifocal to diffuse increase in interstitial cellularity, mild to moderate mononuclear vasculitis with lymphocyte dominated perivascular infiltrates and multiple larger focal areas with occasional degenerate epithelial cells and some infiltrating lymphocytes, with viral antigen expression in both type I and II pneumocytes (Supplemental Fig. S2). Omicron infected mice exhibited only mild pulmonary changes, with mild patchy increase in interstitial cellularity, mild perivascular lymphocyte infiltrate and several disseminated patches of unaltered appearing alveoli with antigen-positive pneumocytes. At both 6 dpi (studied in infections with all variants) and, less severely, 7 dpi (studied in Pango lineage B, Delta and Omicron infected mice), the lungs of Pango lineage B, Alpha, Beta and Delta variant infected animals exhibited changes similar to those described for day 5 post infection (Supplemental Fig. S2); these have been reported previously in more detail [43, 49–50]. Viral antigen expression was restricted to type I and II pneumocytes of unaltered appearing alveoli and in some infiltrating macrophages. In Omicron infected animals, the lung parenchyma was widely unaltered, though there were focal changes similar to those seen with the other strains; viral antigen expression was observed in the same cell types, though in few and smaller patches of alveoli, as recently reported [49].

At day 5 (studied in Delta and Omicron infected animals), infection of the brain was seen in most Delta variant infected animals, i.e. both mice that had to be sacrificed at this day as they had reached the humane endpoint (20% weight loss), and 4 of the 5 scheduled sacrificed animals. In all positive animals, viral antigen expression was found in numerous patches of neurons in all basal regions, stretching from there towards the cortex. In the Omicron infected mice examined at this time point, there was no evidence of brain infection. Viral antigen expression in the brain was often seen together with its expression in the main olfactory epithelium and the olfactory bulb (Fig. 2A).

**Figure 2.**
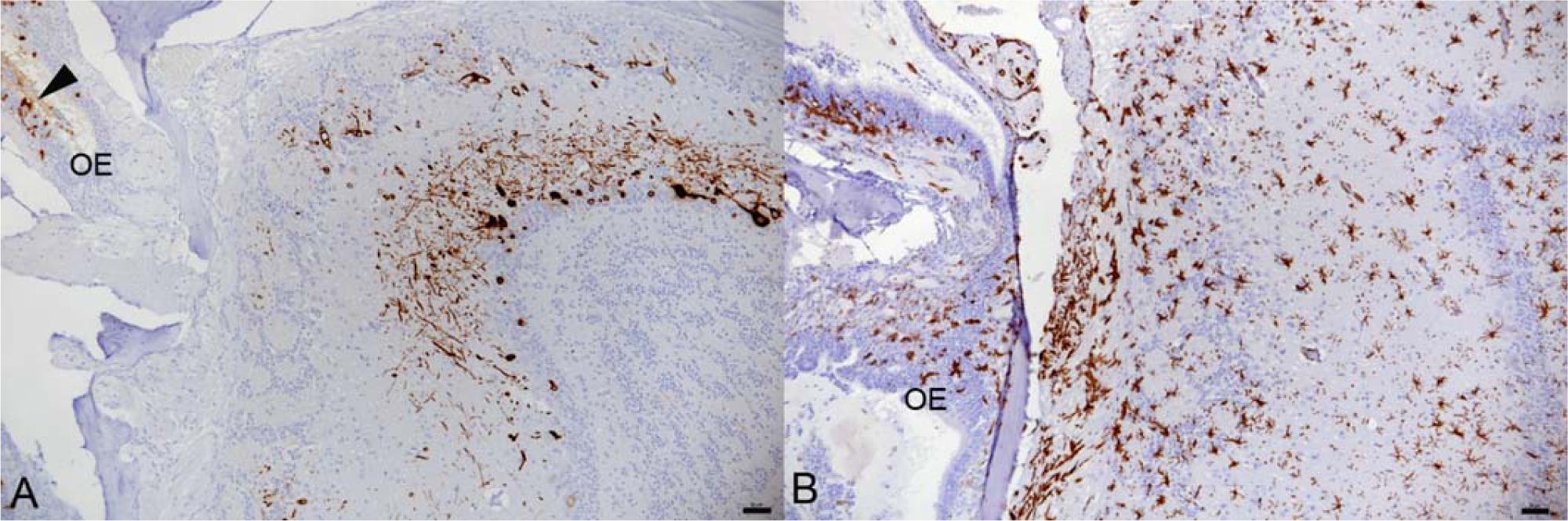
Olfactory bulb, K18-hACE mice, day 5 post intranasal infection with SARS-CoV-2 Delta variant at 10^3^ PFU. Animal with infection of the brain. Consecutive sections showing extensive viral antigen expression in neurons in glomerular, mitral cell and granular cell layer (A) and microglial activation as indicated by the morphology and strong Iba1 expression of microglial cells which is strongest in the olfactory nerve layer (arrow) (B). The olfactory epithelium (OE) exhibits several infected epithelial cells (A, arrowhead) and mild macrophage (Iba1+) infiltration (B).

At day 6, 3 of the 6 Delta variant infected mice exhibited infection of the brain, whereas all 6 Pango lineage B and both Alpha and Beta variant infected mice were found to be positive. At 7 dpi, in Pango lineage B infected animals viral antigen was detected in the brain of 7 of the 9 mice and three of the four IAV co- infected mice (all had received 104 PFU). IAV antigen was not detected in the brain of double infected animals. Of the 6 Delta variant infected mice examined at this time point, 5 were positive in the brain. Again, infection of the brain was often seen together with viral antigen expression in the main olfactory epithelium and the olfactory bulb (Fig. 3A).

**Figure 3.**
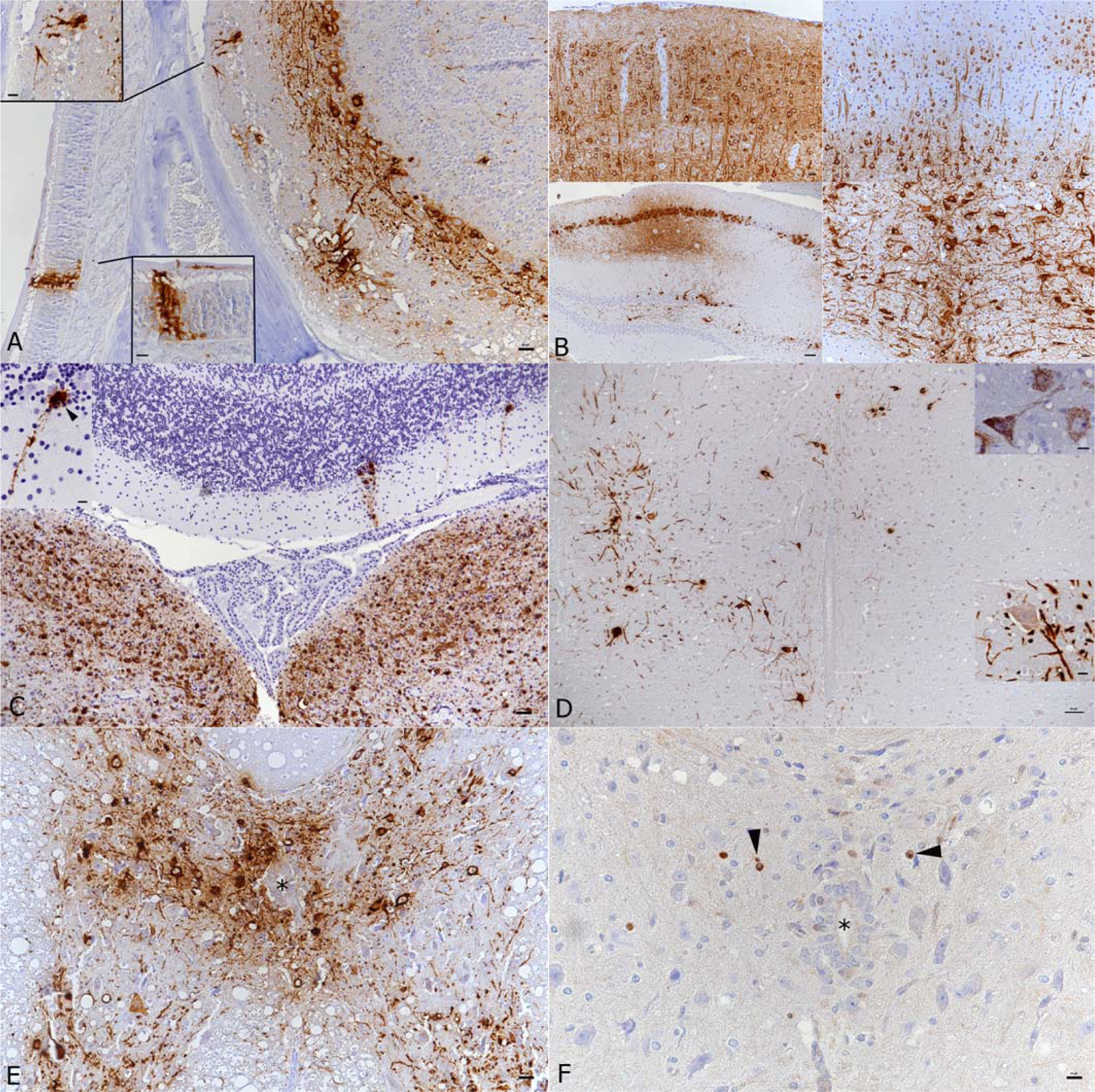
Brain and spinal cord, K18-hACE mice, day 7 post intranasal SARS-CoV-2 Pango lineage B infection (10^4^ PFU as single infection, or as double infection after initial infection with IAV (10^2^ PFU IAV strain A/X31)). A) Double infected animal, main olfactory epithelium (MOI), cribriform plate and olfactory bulb. Viral antigen is detected in olfactory neurons and basal cells of the MOI and in granule layer, inner and outer plexiform layer, mitral layer as well as glomerular layer of the olfactory bulb. Bar = 20 µm. B) Double infected animal, examples of viral antigen expression in different brain regions. Top: frontal cortex with viral antigen expression in almost all neurons (left) and mid cortex with proportion of positive neurons (right); bar = 20 µm. Bottom: patchy virus antigen expression in the hippocampus (CA1 and CA3; left; bar = 50 µm) and strong expression in the medulla oblongata (right; bar = 20 µm). C) Single infected animal, medulla oblongata and cerebellum. Viral RNA is abundantly expressed in neurons in the medulla oblongata (vestibular nuclei). The cerebellar cortex exhibits a few positive Purkinje cells (see al-so inset). Bar = 50 µm. D) Single infected animal, thoracic spinal cord. The grey matter exhibits numerous neurons that express viral antigen (large image and bottom inset) and viral RNA (top inset) in cell body and processes. Bar = 50 µm. E F) Double infected animal, thoracic spinal cord. There is extensive viral antigen expression in neurons in the grey matter (E). Consecutive section showing scattered apoptotic (cleaved caspase 3 positive) glial cells (F, arrowheads), among intact neurons and in the absence of an inflammatory reaction. * - central canal. Bars = 20 m. Immunohistochemistry and RNA-ISH, hematoxylin counterstain.

In none of the Omicron infected animals was there any evidence of viral antigen expression in brain nerves and brain including olfactory bulb at 5, 6 or 7 dpi.

In K18-hACE2 mice, SARS-CoV-2 infection in the central nervous system is restricted to neurons and spreads in the brain and into the spinal cord In the present study, all animals that were examined from day 5 post infection onwards found to harbor viral antigen in the brain showed widespread neuronal expression, but with a variable extent and some evidence of a time dependent manner.

The two Delta variant infected mice that had to be sacrificed at day 5 post infection due to humane endpoint (20% weight loss) exhibited patches of positive neurons in olfactory bulb (Fig. 2) as well as all basal regions and stretching from there to the cortex, sparing the cerebellar cortex.

In the IAV and SARS-CoV-2 double infected animals as well as in the higher dose single infected animals, the virus had spread to the spinal cord where it was found in neurons in the grey matter (motor neurons and sensory neurons; Fig. 1D, E), stretching through the entire cervical and thoracic spinal cord, or decreasing progressively from the cervical spinal cord.

RNA-ISH yielded similar results as immunohistology (Fig. 1C), confirming widespread neuronal infection accentuated in the midbrain.

All Pango lineage B, Alpha and Delta variant infected mice found to harbor virus in the brain at 6 dpi exhibited positive neurons in olfactory bulb, cortex, brain stem, hippocampus and medulla. In the Delta variant infected animals, a marked increase in the number of positive neurons was observed in comparison to 5 dpi. Mice infected with the Beta variant showed a higher variability in the number of positive neurons, with patches of positive neurons in basal regions and cerebral cortex sparing the cerebellum.

At 7 dpi, in Delta variant infected mice, the number of positive neurons was slightly lower than at 6 dpi, with a more patchy distribution. In Pango lineage B infected mice, of which a total of 13 infected brains at 7 dpi were included, a strong, nearly diffuse (in affected regions) bilateral immunoreactivity of neurons was detected. In these animals, the extent of infection in the olfactory bulb was investigated more closely.

Expression was variable and ranged from individual to numerous positive neurons (individual cells/neuronal processes in olfactory nerve layer, glomerular layer, almost all cells in external plexiform layer, individual cells in mitral cell layer), and sometimes also aggregates of positive sensory neuronal cells and their dendrites in the olfactory epithelium layer of the main olfactory epithelium (MOE; Fig. 3A). None of the animals exhibited a reaction in cranial nerves or the inner ear.

Overall, viral antigen was detected in a widespread manner across most brain regions in all infected animals (Fig. 3B, Supplemental Fig. S2). Positive neurons were found among others in anterior olfactory nucleus, primary and secondary motor area, primary somatosensory area, anterior cingulate area, gustatory area, auditory area, infralimbic area, lateral and medial septal nucleus, caudoputamen, piriform area, visual area, ectorhinal area, entorhinal area, retrosplenial area, hippocampus and dentate gyrus, subiculum, nearly all midbrain nuclei, thalamus/hypothalamus, amygdalar nuclei, nucleus accumbens, several cranial nerve nuclei (trigeminal, vestibular, hypoglossal nuclei), reticular nucleus, cuneate nucleus, dentate nucleus. These areas are depicted in a 3D reference atlas of the Allen Mouse Brain Common Coordinate Framework [53; open access] in the following link: https://connectivity.brain-map.org/3d-viewer?v=1&types=PLY&PLY=453%2C1057%2C31%2C44%2C254%2C500%2C922%2C895%2C549%2C1097%2C386%2C370%2C1132%2C987%2C669%2C1089%2C672%2C502%2C909%2C961%2C972%2C507. Viral antigen expression was most intense in the midbrain. The cerebellum showed only individual or small groups of SARS-CoV-2 positive neurons in a few individual animals (Fig. 3C).

Pango lineage B and IAV double infected animals displayed a similar infection of neurons in nearly all brain regions; the expression appeared overall more widespread than in single infected animals.

We also examined the cervical and thoracic spinal cord in the Pango lineage B single and IAV double infected animals. Indeed, the virus had spread to the spinal cord where it was found in neurons in the grey matter (motor neurons and sensory neurons; Fig. 3D, E), stretching through the entire cervical and thoracic spinal cord, or decreasing progressively from the cervical spinal cord.

RNA-ISH yielded similar results as immunohistology (Fig. 3C), confirming wide- spread neuronal infection accentuated in the midbrain.

### SARS-CoV-2 infection of the brain is associated with diffuse microglial activation and a mild macrophage and T cell dominated inflammatory response

In animals where virus was detected in neurons in the brain at 3 and 5 dpi of Pango lineage B and at 5 dpi of the Delta variant infection, this was not associated with any histopathological changes. However, infection in the olfactory bulb was found to be associated with diffuse microglial activation in the area (Fig. 2B). At 6 and 7 days post Pango lineage B infection, a mild nonsuppurative (meningo)encephalitis was consistently observed. The inflammation was only present in areas with SARS-CoV-2 infected neurons. Therefore, it was most pronounced in frontal coronary sections of caudoputamen and the thalamus/hypothalamus region as well as in the hippocampal area where it was represented by infiltration of the wall and the perivascular space of small veins by predominantly mononuclear cells, accompanied by a mild increase in parenchymal cellularity (Fig. 4A). The majority of cells in the (peri)vascular infiltrates were Iba1-positive macro-phages (Fig. 4B). These were accompanied by T cells (CD3+) which comprised up to approximately 30% of the perivascular cells (Fig. 4C). Individual T cells were also found in the neuroparenchyma, mostly in proximity to perivascular infiltrates (Fig. 4C). The vascular infiltrates contained rare individual neutrophils (Ly6G+) and B cells (CD45R/B220+); Fig. 4D). Animals infected with Alpha (6 dpi), Beta (6 dpi) and Delta (6 and 7 dpi) variants and also the Pango lineage B and IAV double infected animals exhibited a similar inflammatory reaction,both in extent and composition. Omicron infected with (all negative for viral antigen) did not exhibit any histological changes in the brain.

**Figure 4.**
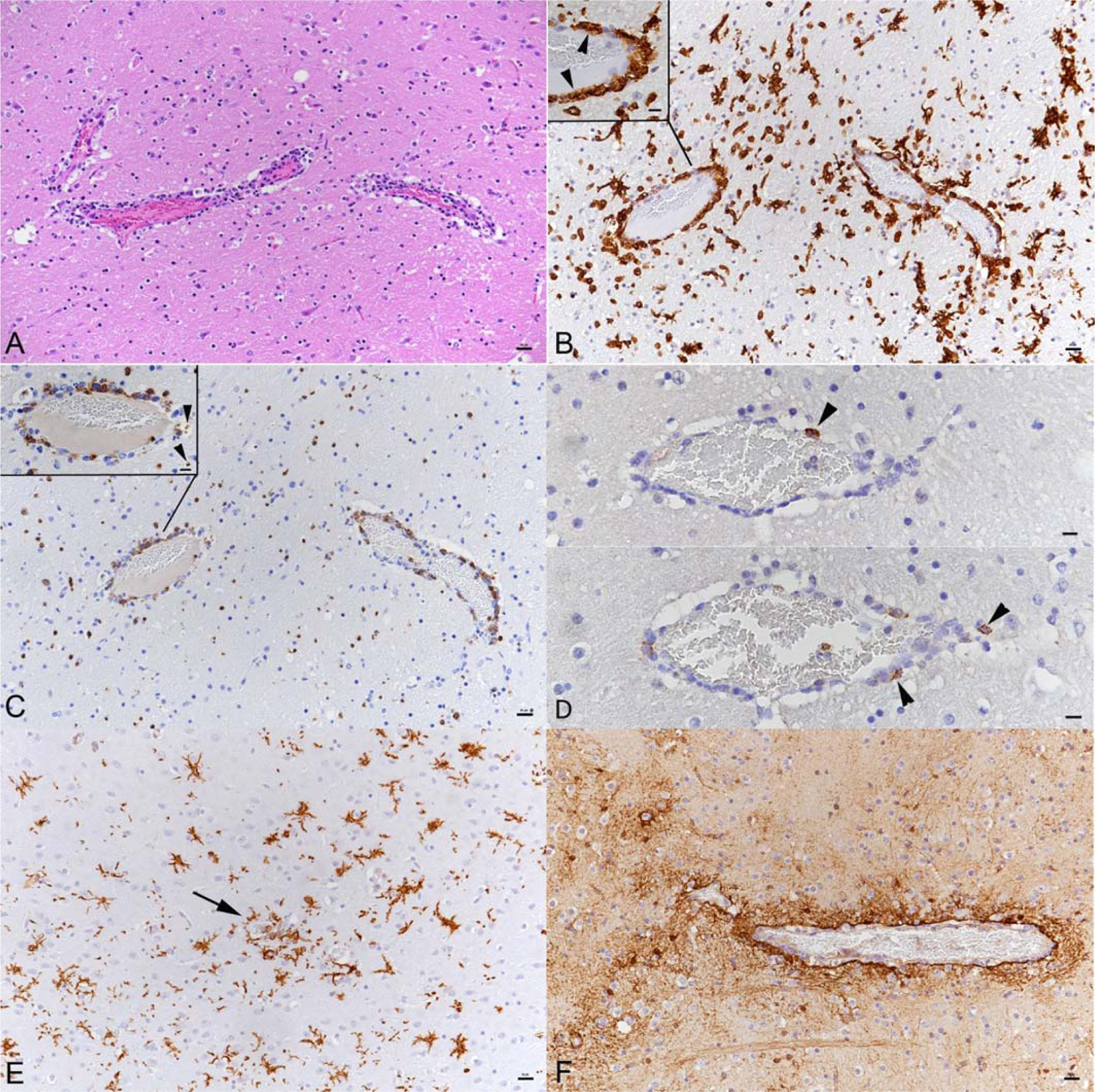
Brain, K18-hACE mouse, day 7 post intranasal SARS-CoV-2 Pango lineage B infection (104 PFU) after initial infection with IAV (10^2^ PFU IAV strain A/X31). A) Brain stem with moderate (peri)vascular leukocyte infiltration and mildly increased cellularity in the parenchyma. HE stain. B) Macrophages (Iba1+) dominate in the (peri)vascular infiltrate (inset: arrowheads) and the surrounding parenchyma exhibits microglial activation (Iba1+ stellate shaped cells). C) T cells (CD3+) are also abundant in the (peri)vascular infiltrates and are found infiltrating the adjacent parenchyma. Inset: vessel with infiltrating T cells; some are degenerate (arrowheads). D) Neutrophils (Ly6G+; top, arrowhead) and B cells (CD45R/B220+; bottom, arrowheads) are very rare in the infiltrates. E) Brain stem with activated microglial cells (Iba1+) and small microglial nodule (arrow). F) Staining of astrocytes (GFAP+) shows hypertrophied astrocytes around a vessel with a mild perivascular infiltrate. B-F: immunohistochemistry, hematoxylin counterstain. Bars = 20 μm.

The inflammatory infiltrates were accompanied by moderate diffuse microglial activation/microgliosis in areas with SARS-CoV-2 infected neurons (Figs. 4B, E and 5A, B). Microglial nodules were detected mainly adjacent to areas with perivascular infiltrates (Fig. 4E). A similar extent of microglial activation and microgliosis was detected at both days 6 and 7 post infection with any of the variants. In the Delta variant infected mice with brain involvement, infected neurons were accompanied by mild microglial activation and microgliosis in close proximity (Fig. 2A, B). Mice infected with the Omicron variant did not show any evidence of microglial activation or microgliosis.

There was no evidence of prominent astrogliosis. However, astrocytes immediately adjacent to perivascular infiltrates showed hypertrophy of the cytoplasm consistent with activation (Fig. 4F).

There is no evidence of viral infection of glial cells, endothelial cells or leukocytes or of neuronal cell death, axonal damage or demyelination in association with SARS-CoV-2 infection, but of apoptotic death of endothelial cells and immune cells.

Despite widespread infection of neurons, there was no morphological evidence of neuronal cell death. Staining for cleaved caspase 3 did not mark any neurons. Also, staining for amyloid precursor protein (APP) to detect disturbances in fast axonal transport as an indicator of acute axonal damage did not show any evidence of the latter. Furthermore, there were no changes indicating obvious demyelination of the white matter. Nevertheless, bilateral and symmetrical myelin sheath vacuolation was detected in the white matter tracts of the pons and the brainstem of animals at 3 and 7 dpi with the Pango lineage B strain, both at low and high doses. Furthermore, mild to moderate vacuolation of the cerebellar white matter without any cellular infiltrates or gliosis was observed. The vacuolation was also present to a mild degree in the animals infected with the Beta strain at 3 dpi, with the Delta strain at 5, 6 and 7 dpi. Capillary endothelial cells which are posi-tive for ACE2 throughout the brain and spinal cord (Fig. 5C) were negative in both SARS-CoV-2 IH and ISH. However, in some vessels that exhibited a leukocyte infiltrate, the endothelial layer appeared focally discontinuous (Fig. 5D), confirming vasculitis. The infiltrate contained scattered degenerate cells suggesting a leukocytoclastic component. (Fig. 5D, E).

**Figure 5.**
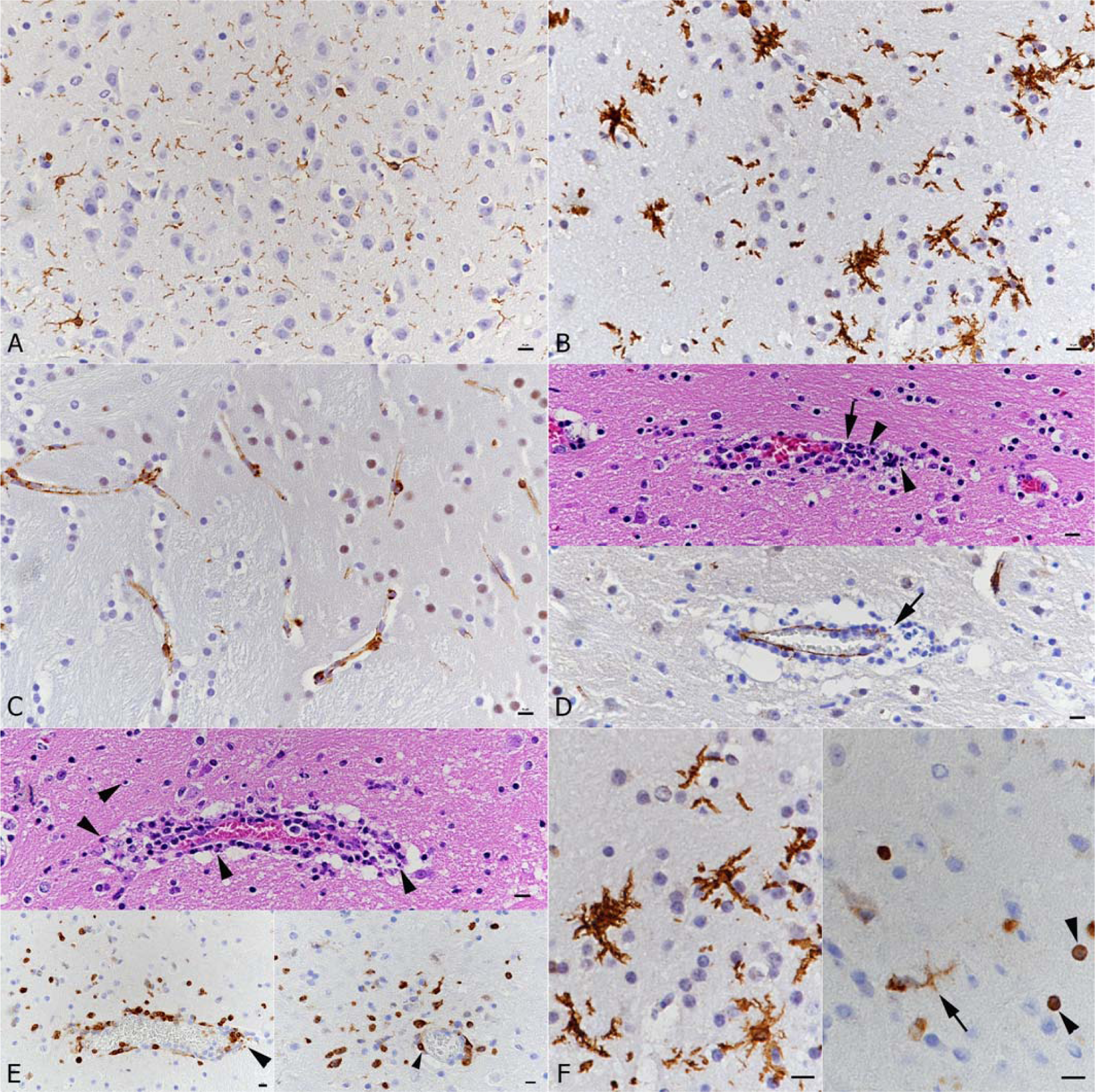
Brain, K18-hACE mice, mock infected and at day 7 post intranasal SARS-CoV-2 Pango lineage B infection (10^4^ PFU) after initial infection with IAV (102 PFU IAV strain A/X31). A) Mock infected mouse. Quiescent microglia (Iba1+). B) Infected animal. Diffuse microglial activation (Iba1+ stellate cells). C) Mock infected mouse. Endothelial cells show cytoplasmic ACE2 expression. D-F) Infected animal. D) Vessel with moderate leukocyte infiltration with focal destruction of the vessel wall (arrows) and some degenerate leukocytes (leukocytoclastic component). Top: HE stain; bottom: staining for ACE2, showing disruption of the endothelial cells layer. E) Vessel with moderate leukocyte infiltration (top: HE stain; bottom: staining for cleaved caspase 3). Individual cells in the infiltrate and in the adjacent parenchyma (top and bottom left: arrowheads) and endothelial cells in an affected vessel (bottom right: arrowhead) undergo apoptosis. F) Activated microglia (Iba1+; left). Staining for cleaved caspase 3 (right) shows apoptosis of a microglial cell (arrow) and of small round cells (arrowheads; morphology consistent with lymphocytes). A-E: bars =20 μm; F: bars = 10 μm.

This was confirmed by staining for cleaved caspase 3; both endothelial cells and leukocytes in the vascular infiltrates were found to be apoptotic. Apoptosis (cleaved caspase 3+) was also observed in a few leukocytes (morphology consistent with lymphocytes) in the adjacent parenchyma (likely infiltrating T cells; Fig. 5E) and in occasional microglial cells (Fig. 5F). In none of the brains was there any evidence of SARS-CoV-2 antigen or RNA expression in glial cells.

## Discussion

COVID-19 is primarily a respiratory disease, with potentially fatal systemic complications that can also involve the CNS. At present, the pathogenetic and clinical role of en-cephalopathy in COVID-19 patients is controversially discussed. A variety of mild and/or severe clinical signs have been described, and the reported morphological findings in the brain differ between studies; similarly, the presence and distribution of viral RNA or anti-gen appears to be inconsistent. Many questions are currently still open regarding the in-fectious route and spatio-temporal distribution of SARS-CoV-2 in the CNS of COVID-19 patients without comorbitidies, confirming the need for appropriate animal models.

In this context, the hACE2 transgenic mouse under the control of the human cytokeratin 18 promoter, a commonly used mouse model to study pathogenetic effects of SARS-CoV-2 infection, has been employed not only in the present, but also in several other recent studies using the USA-WA1 strain, predominantly at high doses (10^5^ to 1.5 x 10^6^ PFU) [26,42,44,54–56]. All these studies, including the present one, show that intranasal inoculation of K18-hACE2 mice with all SARS-CoV-2 variants of concern tested up to now, but not the Omicron variant can lead to infection and extensive virus spread in the brain within 5 to 7 days. Infection even stretches into the spinal cord and can basically affect the entire grey matter, including, though only rarely, also the cerebellar cortex (Purkinje cells). This indicates that SARS-CoV-2 has no selective neurotropism. Interestingly, we found no evidence of brain involvement with Omicron infections which would be consistent with the observation made by this and other groups that these strains are less virulent and result in limited infection in K18-hACE2 mice [57]. Omicron variants have a spike protein that differs substantially from that of other variants [58]. It might bind better to ACE2 but less efficiently to whatever the receptor/counter receptor is in neurons.

In the present study, intranasal inoculation of mice with a low (10^3^ PFU) and higher (10^4^ PFU) viral dose (compared for the Pango lineage B strain) resulted in variable extents of CNS infection and did reach the spinal cord with extensive brain infection. With IAV pre-infection, it was consistently widespread. This does not confirm a direct dose de-pendence of viral spread, different to what was suspected other studies [45, 55], but suggests some effect of prior damage in the respiratory tract [43]. At the same time, we and others found virus in the olfactory epithelium and in neurons in the olfactory bulb [42,44–45]. Therefore, a rostral to caudal spread of infection with emphasis on basal structures and consequent infection of cortical areas showing a more patchy than diffuse pattern is most likely. This implies a direct neuronal spread of the virus rather than a hematogenous route of infection which would likely result in a more disseminated infection pattern as suggested also by other authors [8]. This might also explain the spatio-temporal antigen distribution in Delta variant infected mice with an increase from 5 to 6 dpi and a decrease in positive neurons from 6 to 7 dpi as well as a more patchy than diffuse distribution pattern. An ultrastructural examination on SARS-CoV-2 infected human brain organoids is in agreement with the theory of neuronal spread of the virus, as it provides evidence of virus cell-to-cell-spread between neurons in the organoids [26]. Many of the SARS-CoV-2 antigen positive neurons of mice of this study are located in areas which are secondary or tertiary connections of the olfactory bulb. This gives further evidence of virus entry via the olfactory bulb, as confirmed in both mice [42, 56] and humans [10] as a natural route of brain infection by SARS-CoV-2. However, many virus-positive regions that are not directly connected to the olfactory system were also identified, providing further evidence of an additional route of viral dissemination [56].

Clinical signs in mice consistent with neurological disease have been quite rarely re-ported, where mice had shown hunchbacked posture, ruffled fur, tremors and ataxic gait from day 4 pi, and died after day 6. These mice carried very high levels of infectious virus in the brain and showed an encephalitis with perivascular hemorrhage. Neuronal death via apoptosis was suspected [44]. Other studies reported perivascular cuffs or vasculitis, neuronal degeneration and necrosis, satellitosis, parenchymal edema, and occasional mi-crothrombi [42,45,54–56], all at day 7 pi. The present study confirms that SARS-CoV-2 variants readily infect neurons in the K18-hACE2 mice, with viral protein accumulation in the entire cytoplasm including the cell processes but does not provide evidence of neu-ronal cell death in association with infection, indicating that SARS-CoV-2 has no direct cytopathic effect on neurons. This may be due to the lower viral dose, provided that a high viral load would damage neurons directly.

Involvement of the spinal cord, which was a prominent feature in our mouse model, has only been reported in a few human cases [59]. It might be a so far under- recognized neurological complication, since neuropathological assessments did not go beyond neuroimaging and evaluation of several CSF parameters.

Similar to Carossino et al. [42], the present study did not find evidence of demyelination. Clusters of swollen neurons with foamy or vacuolated cytoplasm, occasionally with pyknotic and eccentric nuclei, as observed by [45] were not detected in our mice. Furthermore, axonal damage was not present in the investigated murine brain sections. However, while both axonal damage and demyelination can occur in COVID-19 patients, they are apparently not common neuropathological findings [18-19,21]. Similar to Vidal et al. [45], we also observed mild spongiosis of tracts of the medulla oblongata in some in-fected mice, although axons in these tracts were SARS-CoV-2 antigen negative. The relevance of the vacuolation seen in brain stem and cerebellar white matter which was most evident in animals infected with the Pango lineage B strain remains questionable. It also needs to be considered that vacuolation, affecting principally white matter can be a consequence of prolonged holding of formalin fixed tissue in 70 % alcohol, as shown in bo-vine brains submitted for diagnostic histopathology [60]. This approach, mainly taken to preserve immunoreactivity when processing of the tissue is delayed, was also chosen for the present study; it has the additional advantage that it allows the shipping of tissue which formalin does not. Spongiosis as a common artifact was also described by Wohlsein et al. [61] discussing that it is difficult to discriminate between early significant and non-significant changes.

Neuronal SARS-CoV-2 infection without obvious neuronal degeneration is reminiscent of other neurotropic and persistent viral infections, like herpes simplex virus (HSV). HSV-1 establishes latency in the mouse model within the earliest stages of acute infection, with the viral genome reaching the neuronal ganglia within the first 24 hours of infection [62]. Further studies would now be required to determine whether SARS-CoV-2 also establishes latency or whether there is subsequent neuronal damage.

Viral infection of neurons is evidently associated with an antiviral response in the mice, as indicated by a reported increase in IFN-α mRNA in the brain from day 5 pi onwards, with a peak of both mRNA and protein on day 6 [44]. This is accompanied by an inflammatory response in the brain parenchyma. Similar to a previous study [54], we observed a mild non-suppurative, macrophage and T cell driven encephalitis (perivascular infiltrates and occasional vasculitis) in all mice that harbored viral antigen in the brains at 6 and 7 dpi. This appeared to be slightly more intense with more extensive neuronal virus antigen expression in animals that had been preinfected with IAV, suggesting easier access to the brain with pre-existing damage due to IAV infection [43]. Transcriptional investigations also confirmed the inflammatory state of the brain, demonstrating an increase in inflammatory cytokines and chemokines (IL-6, TNF-α, IFN-γ, IL-1β, MIP-1α, MIP-2, IP-10) in the brain of infected mice [44, 55]. This might also explain the activation/hypertrophy of astrocytes in areas with perivascular infiltrates and the diffuse microglial activation and multifocal microgliosis that we and others observed [42,55,63]. Both glial cell populations are responsive to proinflammatory signals from endothelial cells, macrophages and/or neurons [6]. The proinflammatory gene expression program initiated after viral infection would expand neuroinflammation [28]. Interestingly, microgliosis was also shown in the olfactory bulb of infected mice without obvious inflammatory cell infiltration similar to human patients [20]. Activated microglia may itself undergo a phenotypic shift and display exaggerated release of proinflammatory mediators and aberrant phagocytic activity, inducing neurodegeneration [64]. This could explain the diffuse astrogliosis reported from other mouse and human studies [9,42,45] as well as neuronophagia by microglial cells which was seen in some human COVID-19 patients [23] and one mouse study [42].

In previous studies we and others have shown that IAV pre-infection of K18- hACE2 mice results in more severe SARS-CoV-2 associated pulmonary changes and higher amounts of infectious virus at 6 or 7 days post Pango lineage B infection [43, 65]. Interestingly though, viral loads in the brain were not found to be increased [65], although we observed more widespread brain infection in animals that were infected with IAV. It is currently unclear which direct impact IAV infection has for the subsequent SARS-CoV-2 challenge, but more efficient virus entry could be an option, since IAV infection was shown to upregulate ACE2 expression in vitro [65]. In co-infected hamsters, an involvement of IL-6 in the increased severity of pneumonia was discussed [66].

While we did not find evidence of neuronal death, we observed apoptosis of infiltrat-ing lymphocytes, capillary endothelial cells and macrophages/microglial cells in proximity to perivascular infiltrates. As in a previous study [42], none of these cells were found to be SARS-CoV-2 infected. This result is in line with recent findings in SARS-CoV-2 infected human brain organoids which provided evidence that infected neurons do not die but can promote the death of adjacent uninfected cells [26]. Using single cell RNA sequencing, the study showed that SARS-CoV-2 infected cells in the organoids were in a hypermetabolic state, whereas uninfected cells nearby were in a catabolic state and a hypoxic environment, as shown by HIF1α expression [26]. The same may be true for the brain of K18-hACE2 mice with widespread neuronal SARS-CoV-2 infection. Also, coupled proliferation and apoptosis are assumed to maintain the quite rapid turnover of microglia in the adult brain [67]. We did not determine the proliferation rate in this study, but such “physiological” turnover should be considered when interpreting microglial apoptosis in SARS-CoV-2 infected mice. In another hACE2 mouse study, with prominent microglial activation, pro-inflammatory CSF cytokines/chemokines were elevated for at least 7 weeks post infection, a feature also reported in human patients with long COVID syndrome [68]. Also in vitro neuronal cell cultures of K18-hACE2 mouse neurons showed an upregulation of the expression of genes involved in innate immunity and inflammation [69]. An astrocytic SARS-CoV-2 infection which was described in human brains and cultured astrocytes [36] could not be detected in any mouse brain with any virus variant in our study.

Therefore, the present study found no evidence that SARS-CoV-2 infects astrocytes. Nevertheless, due to the controversial findings further studies are needed to elucidate the role of astrocytes in SARS-CoV-2 infection. Several morphological changes in the mice used in the present study recapitulate findings reported in brains of human COVID-19 patients, like mild perivascular inflammatory infiltrates [21] and microgliosis [20]. Further vascular lesions apart from vasculitis/endotheliitis, like ischemic infarcts [18, 26] or microthrombi [16] that seem to be fre-quent in fatal human COVID-19 cases, are obviously not a regular feature in this mouse model, since only two studies reported occasional microthrombi in the brain [55–56]. This could be due to the lack of endothelial cell infection in the brain of the mice, whereas it was seen in association with fresh ischemic infarcts in the brain of a COVID-19 patient [10, 26]. In our study, mice infected with the delta variant did not show inflammatory changes but needed to be euthanized due to clinical symptoms. Interestingly, the presence of virus in the brain of the human patients is apparently also not consistently associated with leukocyte infiltration, indicating that SARS-CoV-2 does not necessarily induce an immune response like other neurotropic viruses [26].

The main host receptor for SARS-CoV-2 is the human angiotensin-converting enzyme 2 (hACE2). While a study in the brain of fatal human COVID-19 cases showed ACE2 expression in cortical neurons and found evidence that ACE2 is required for infection of human brain organoids [26], neuroinvasion and spread in the K18-hACE2 mice is apparently not directly dependent upon ACE2 expression, because the virus does not infect all ACE2 expressing cells and does infect cells without apparent ACE2 expression [42]. Indeed, in line with the findings of previous studies we only detected ACE2 protein expression in capillary endothelial cells, ependymal cells and choroid plexus epithelium in the brain and spinal cord, in the absence of detectable viral antigen [26, 42–43]. In human COVID-19 patients, viral particles were detected in the endothelium of capillaries in kidney, liver, heart, lung and small intestine. This was associated with an endotheliitis [70]. This finding was discussed controversially [14]. Furthermore, ultrastructural examination of the brain of one human patient showed viral particles in small vesicles of endothelial cells, suspecting an additional hematogenous route of brain infection [12]. Indeed, using intravenously injected radioiodinated SARS-CoV-2 S1 it has been shown that the viral protein can cross the BBB, and likely by absorptive transcytosis [34]. It may hence be due to the fact that endothelial ACE2 expression in the K18-hACE2 mice represents only the murine protein [42] that the brain endothelium of the mice does not become infected by SARS-CoV-2. The role of other receptors is still subject of debate, a possible candidate might be neuropilin-1 [8].

Based on current knowledge, it seems likely that in COVID-19 patients the virus can access the brain in several ways. In severe COVID-19, viremia could allow access to the brain via the vasculature [71]. With the platelet hyperactivity in critically ill patients [72] and abnormal blood clotting or unusual thrombotic presentations [73–74] this might lead to the ischemic changes described in a proportion of fatal cases, without viral infection of neurons. This might not be mirrored by the murine model as viremia may not occur [55]. In mild COVID-19, i.e. without viremia, infection of the brain via the olfactory bulb and possible other neurogenic routes could occur. This would then lead to exclusive neuronal infection and an only mild inflammatory response. It might be this scenario that explains the loss of taste and smell and severe headaches reported in patients with mild disease.

The murine model does not fulfil all morphological criteria of acute human neurological COVID-19. However, brain infection following a lower dose of intranasal challenge might represent a mouse model for long COVID-19 syndrome studies. The pathogenesis of this sequel of the acute disease is still unknown, but fatigue, muscle aches, breathlessness, and headaches are the most frequently reported symptoms [75–76]. A persistent low-grade smoldering inflammatory response to newly budding virions might be central to the con-dition. Also, degeneration or impaired function of neuronal and glial cells that are cardi-nal for the physiological function of the brain might be an option [77]. A mild inadequate immune response with persistent viral load and viral evasion of the immune surveillance are suspected key factors [77].

This is supported by the fact that COVID-19 patients still exhibit a significant remaining inflammatory response in the serum at 40-60 days post infection [78]. Further investigations of the inflammatory response at transcriptome and translational level in the murine model could provide further insights.

These data also have important implications for the development of therapeutic interventions. Firstly, the endothelium and astrocyte foot processes represent key compo-nents of the BBB, protecting the brain from accumulation of endo- and xenobiotics. Future studies should address the consequences of infection for maintenance of barrier integrity to mitigate potential inadvertent delivery of neurotoxic agents that would otherwise not permeate the brain. Secondly, several studies have already sought to understand pulmonary distribution of postulated therapeutic interventions [79–81] but robust efficacy of antivirals and/or immunomodulatory agents may also necessitate adequate exposure within the CNS. Notably, the repurposed antivirals remdesivir, favipiravir and molnupiravir exhibit low concentrations in the brain relative to plasma in preclinical species [82–85]. Furthermore, dexamethasone also exhibits low brain penetration in mice with an intact BBB, but this increases in mice genetically engineered to be absent a critical drug transporter, P-glycoprotein [86].

Further work is required to define the importance of brain penetration of therapeutics being investigated as interventions across the spectrum of disease from prevention, mild, moderate, severe to long COVID-19.

## Conclusions

Despite widespread neuronal infection, pathomorphological changes of the course of disease consisted of mild lymphohistiocytic inflammation and microglial activation.

Solely neuronal cells were infected which supported the infectious route via the olfactory epithelium, olfactory bulb and transsynaptic spreading. The limited expression of ACE2 raises the question for ACE2-independent pathogenetic mechanisms to explain the neurotropism of the virus. Microgliosis and immune cell apoptosis were main pathological features in our study indicating a potential important role of microglial cells in the pathogenesis of neuromanifestation in COVID- 19.

## Acknowledgements

We are grateful to the technical staff in the Histology Laboratory, Institute of Veterinary Pathology, Vetsuisse Faculty, University of Zurich (IVPZ), for excellent technical support. This work was funded in part by the US Food and Drug Administration (USA) 75F40120C00085, Characterization of severe coronavirus infection in humans and model systems for medical countermeasure development and evaluation. Work in the lab is also supported by MRC grant MR/W005611/1, G2P-UK; A National Virology Consortium to address phenotypic consequences of SARSCoV-2 genomic variation.

## CONFLICT OF INTEREST

The authors declare that they have no conflict of interest.

## Notes

### Competing Interest Statement

The authors have declared no competing interest.

### Summary of Updates

The manuscript has been revised after we extended the study by investigating mice infected with other virus variants.

