## Supplementry Table and Figure for "Neuroinvasion and neurotropism by SARS-CoV-2 variants in the K18-hACE2 mouse"

**Supplemental Material**

**Supplemental Table S1**. Antibodies and detection methods used for immunohistochemistry

| **Antigen** | **Antibody (clone)** | **Antigen retrieval** | **Antibody dilution, detection method** |
| --- | --- | --- | --- |
| SARS-CoV NP | pAB rabbit^a^ | CB pH 6.0,  98 °C, 20 min | 1:6,000, ON, 4 °C  EnVision+ (HRP, Rb)^f^ |
| IAV | goat anti-IAV (H1N1; virions)^b^ | TE pH 9.0,  96 °C, 30 min | 1:200, 60 min, RT  Rb anti-goat Ig/HRP^f^ |
| Iba1 | pAB rabbit anti-human Iba1^c^ | CB pH 6.0,  98 °C, 20 min | 1:750, 60 min, RT  HRP EnVision+ (HRP, Rb)^f^ |
| CD3 | mAB rabbit anti-mouse CD3 (SP7)^d^ | TE pH 9.0,  37 °C, 20 min | 1:900, 60 min, 37 °C, OmniMap anti-Rb HRP^i^ |
| CD45R | mAB rat anti-mouse CD45R (B220/RA3-6B2)^e^ | CB pH 6.0,  98 °C, 20 min | 1:800, 60 min, RT  EnVision+ (HRP, Rat)^f^ |
| GFAP | pAB rabbit anti-bovine GFAP^f^ | CB pH 6.0,  98 °C, 20 min | 1:600, 10 min, RT  MACH4™ HRP^k^ |
| APP | mAB mouse anti-Alzheimer precursor A4 (pre A4695) fusion protein (22C11)^g^ | CB pH 6.0,  98 °C, 20 min | 1:6,000, 60 min, RT  MACH4™ HRP^k^ |
| ACE2 | mAB rabbit anti-human ACE2 (SN0754)^h^ | CB pH 6.0  100 °C, 20 min | 1:200, 10 min, RT  EnVision+ (HRP, Rb)^f^ |
| Cleaved caspase 3 | mAB rabbit anti-cleaved caspase 3 (Asp175, 5A1E)^l^ | TE pH 9.0,  98 °C, 20 min | 1:200, ON, 4 °C  EnVision+ (HRP, Rb)^f^ |

**Legend**: ACE2 - angiotensin-converting enzyme 2, APP - Alzheimer Precursor Protein A4, GFAP – glial fibrillary acidic protein, IAV – Influenza A virus, mAB – monoclonal antibody; pAB – polyclonal antibody; CB – citrate buffer; ON – overnight; RT – room temperature; Rb – rabbit; SARS-CoV NP – severe acute respiratory syndrome coronavirus nucleocapsid protein; TE – Tris-EDTA buffer;

**Commercial providers**:

^a^Rockland Immunochemicals Inc., Limerick, USA

^b^Meridian Life Sciences Inc., Memphis, USA

^c^WAKO, Osaka, Japan

^d^Spring Bioscience Corp., Ventana Medical Systems, Tucson, USA

^e^BD Pharmingen, Franklin Lakes, USA

^f^Agilent Dako, Glastrup, Denmark

^g^Merck Millipore

^h^Novus Biologicals, Centennial, USA

^i^Ventana Medical Systems Inc., Tucson, USA

^k^Biocare Medical

^l^Cell Signaling Technology

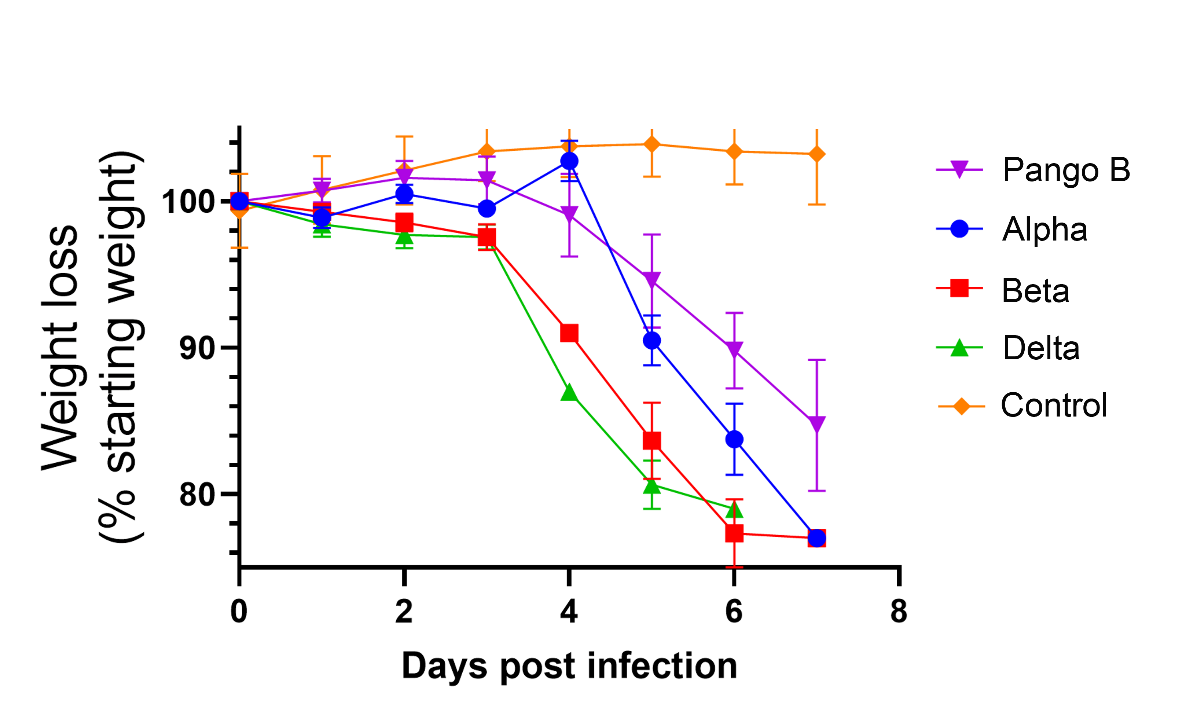

**Figure S1**. **Infection of K18-hACE2 mice with SARS-CoV-2 VOCs leads to substantial weight loss.** K18-hACE2 mice were infected intranasally with SARS-CoV-2 Alpha, Beta or Delta variant at 10^3^ PFU (n=4 per group). Mice were monitored for weight loss at indicated time points and euthanized at days 5 and 6 post infection due to substantial weight loss (limit: 20% loss) or at the end of the experiment (day 7). The weights of Pango lineage B infected animals (10^4^ PFU) from another experiment served for comparison. Data represent the mean value ± SEM.

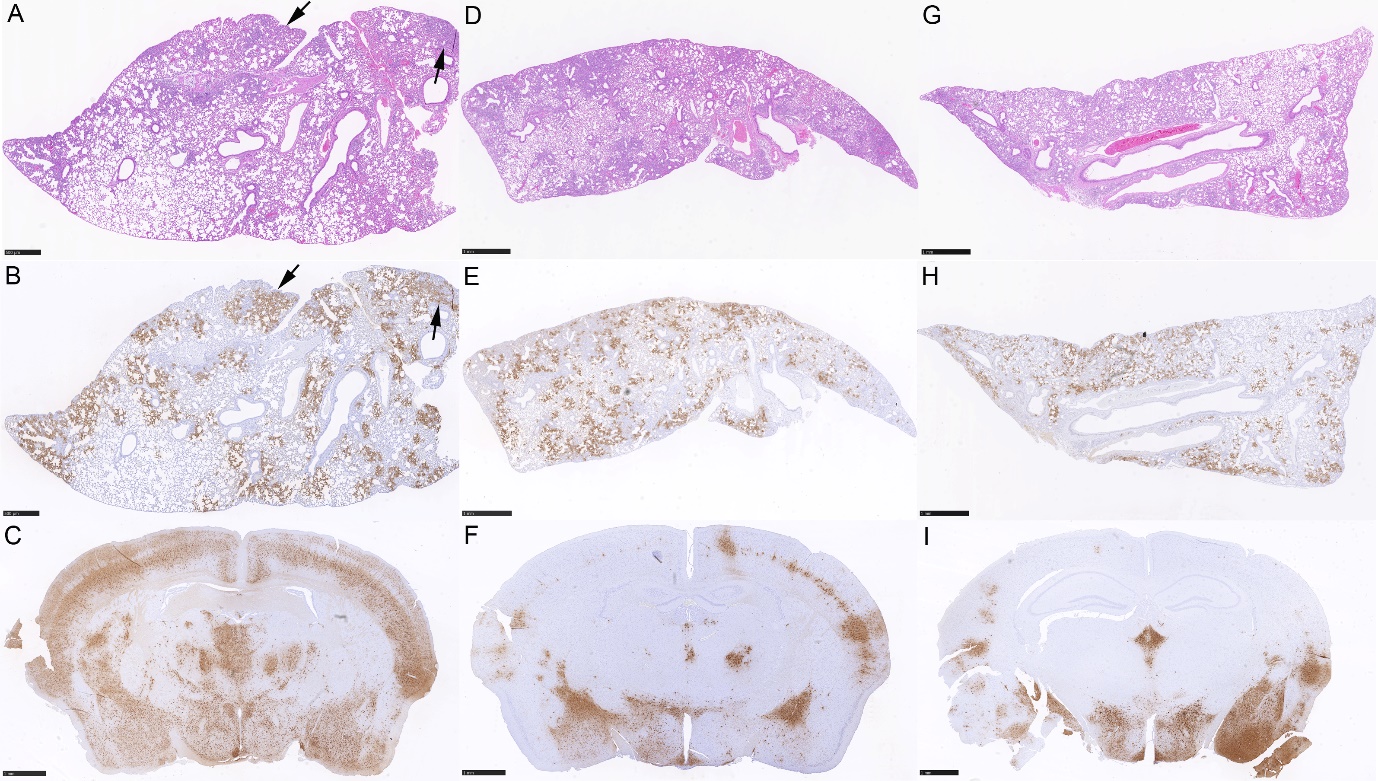

**Figure S2**. **Infection of K18-hACE2 mice with SARS-CoV-2 VOCs leads to extensive pulmonary infection and spreads to the brain.** K18-hACE2 mice were infected intranasally with SARS-CoV-2 Alpha, Beta or Delta variant at 10^3^ PFU and euthanized at 5 or 6 days post infection (dpi) due to substantial weight loss (limit: 20% loss). Lungs and brains, overview. **A-C)** Animal infected with Alpha variant and euthanized at 6 dpi. **A, B)** Lung. **A)** Mild multifocal increase in interstitial cellularity (arrows) with activation of type II pneumocytes and occasional degenerate and sloughed off alveolar epithelial cells. **B)** There is extensive viral antigen expression in large patches of unaltered appearing alveoli and in more cell rich areas (arrows). **C)** Brain, coronal section at level of corpus callosum, caudoputamen, thalamus/hypothalamus and frontal cerebral cortex. There is extensive viral antigen expression in neurons in cerebral cortex, thalamic and hypothalamic nuclei as well as piriform area. **D-F)** Animal infected with Beta variant and euthanized at 6 dpi. **D, E)** Lung. **D)** Mild to moderate multifocal increase in interstitial cellularity with activated type II pneumocytes and degenerate and sloughed off alveolar epithelial cells. **E)** There is extensive disseminated viral antigen expression in large patches of unaltered appearing alveoli and in more cell rich areas. **F)** Brain, coronal section at level of corpus callosum, rostral hippocampus, caudoputamen, thalamus/hypothalamus and frontal cerebral cortex. There is multifocal patchy viral antigen expression in neurons in cerebral cortex, amygdala, thalamic/hypothalamic nuclei, piriform area. **G-I)** Animal infected with Delta variant and euthanized at 5 dpi. **G, H)** Lung. **G)** Mild multifocal increase in interstitial cellularity (arrows) with activation of type II pneumocytes and occasional degenerate and sloughed off alveolar epithelial cells. **H)** There is abundant disseminated viral antigen expression in variably sized patches of unaltered appearing alveoli and in more cell rich areas. **I)** Brain, coronal section at corpus callosum, hippocampus, caudoputamen, thalamus/hypothalamus and frontal cerebral cortex. There is multifocal patchy viral antigen expression in neurons in cerebral cortex, amygdala, thalamic/hypothalamic nuclei, piriform area. HE stain (A, D, G). Immunohistochemistry, hematoxylin counterstain (B, C, E, F, H, I).
